# Position-Dependent NMD Generates Diverse Protein Outcomes

**DOI:** 10.64898/2026.08.24.746594

**Authors:** Nusrat Jahan Pinky, Hanae Sato

**Affiliations:** Division of Nano Life Science, Graduate School of Frontier Science Initiative, Kanazawa University, Kakuma-machi, Kanazawa, Ishikawa 920-1192, Japan; WPI Nano Life Science Institute (NanoLSI), Kanazawa University, Kanazawa, Ishikawa, Japan

## Abstract

Nonsense-mediated mRNA decay (NMD) is a translation-dependent mRNA decay pathway triggered by premature termination codons (PTCs). Although NMD is known to eliminate aberrant transcripts, how PTC position influences cell-to-cell heterogeneity in NMD and the resulting protein outputs remains unclear. Here, we used a single-cell NMD analysis system that quantifies cellular variability based on the GFP/mCherry fluorescence ratio. By combining this system with fluorescence-activated cell sorting (FACS), we show that PTC location critically determines not only NMD efficiency and its variability across cells, but also the spectrum of resulting protein products. These include truncated proteins arising from premature termination, full-length proteins generated through translational readthrough, and N-terminally truncated isoforms produced by downstream reinitiation. Our findings reveal that positional and cellular heterogeneity in NMD contribute to proteomic diversity and may underlie the variable phenotypic severity of genetic diseases caused by PTCs. This work establishes a framework for dissecting NMD regulation and its translational consequences.

**Highlights:** Protein-level readouts reflect mechanisms underlying NMD escape

## Introduction

Gene expression faces a fundamental dilemma when translation terminates prematurely: whether aberrant transcripts are eliminated or allowed to produce protein products. This decision is largely governed by nonsense-mediated mRNA decay (NMD), a conserved pathway that selectively targets transcripts containing premature termination codons (PTCs). PTCs are responsible for a substantial fraction of inherited human disorders, accounting for approximately 10–20% of disease-causing mutations ^1,2^. The resulting truncated polypeptides often lack essential functional domains and may exert dominant negative effects ^3–5^. To mitigate these harmful consequences, NMD recognizes and degrades PTC-bearing mRNAs during aberrant translation termination. This surveillance limits aberrant protein accumulation that could otherwise compromise cellular function ^6–9^. Dysregulation of this mechanism has been implicated in a broad spectrum of genetic disorders such as cystic fibrosis, Duchenne muscular dystrophy, and β-thalassemia, where altered NMD efficiency directly shapes disease severity and phenotypic variability ^2,10–12^.

However, NMD is not an absolute process, and a subset of PTC-containing transcripts escapes degradation^37^. The fate of these NMD-escaped mRNAs remains poorly understood, particularly whether they give rise to functional full-length proteins through translational readthrough or instead produce truncated or alternative protein isoforms. Resolving this distinction is critical, as the resulting protein products may directly influence cellular function and disease outcomes.

Although the basic principles governing NMD activation have been extensively characterized, the determinants of NMD efficiency remain incompletely understood. Early studies demonstrated that PTCs located upstream of the normal termination codon are more likely to trigger NMD than those positioned near the 3’ end of the transcript ^13^. This positional effect is further defined by the “50–55 nucleotide rule,” in which stop codons positioned > 50–55 nucleotides upstream of an exon–exon junction efficiently promote NMD ^14–16^. Mechanistically, this positioning effect is linked to exon junction complexes (EJCs), which remain downstream of PTCs and promote recruitment of NMD factors during the pioneer round of translation ^17–19^. In addition, the faux 3’ UTR model proposes that increased distance between a PTC and poly(A)-binding protein (PABP) impairs normal translation termination and facilitates NMD activation ^20–22^. Additional transcript features, such as UPF1 binding, uORFs, coding sequence context, translation efficiency, and local sequence composition, have also been implicated in regulating NMD efficiency ^15,23,24^. Nevertheless, NMD efficiency remains highly variable across transcripts, genes, and cellular contexts ^7,37,27^, suggesting that current models incompletely explain NMD outcomes.

By contrast, the mechanisms underlying NMD escape remain poorly understood, particularly how escaped transcripts contribute to protein production. Distinct protein outcomes may provide clues to the molecular processes that enable transcripts to evade degradation. One such mechanism is translational readthrough, in which ribosomes bypass a PTC and continue translation, representing one possible outcome of NMD escape. Because NMD depends on translation termination and release factor recruitment, readthrough may interfere with NMD activation by favoring near-cognate tRNA incorporation instead of peptide release. Consequently, both readthrough and NMD efficiency are affected by variables such as PTC position, surrounding nucleotide sequence, and the availability of factors that promote readthrough over translation termination ^26–29^.

In addition to translational readthrough, downstream translation reinitiation represents another mechanism that can alter the fate of PTC-containing transcripts. Translation reinitiation provides an additional route by which translation can resume downstream of a premature termination event, typically occurring when translation terminates near the 5′ end of the coding sequence (often within ∼30 amino acids of the start codon). Under these conditions, ribosomes can reinitiate at downstream start codons to generate N-terminally truncated protein isoforms ^30–32^. Reinitiation efficiency depends on features of the short upstream ORF created by premature termination, the distance to downstream initiation codons, and the retention or functional coupling of initiation factors such as eIF4G and eIF3 ^33,34^. Previous studies have shown that downstream translation reinitiation can promote escape from NMD, particularly when the PTC is located close to the translation initiation region ^32,35,36^. These observations suggest that current models primarily explain whether PTC-containing transcripts become targets of NMD. One possible approach to address this question is to examine the protein products generated from NMD-escaped transcripts, as distinct protein outcomes may reflect different mechanisms of escape, including translational readthrough or downstream translation reinitiation. However, transcripts that evade NMD are likely to represent only a minor fraction within a cell population, making their protein products difficult to detect using conventional population-averaged measurements. Approaches capable of selectively enriching NMD-escaped populations may therefore improve sensitivity for detecting protein outputs associated with NMD escape and provide insight into the mechanisms by which transcripts evade degradation.

To overcome these limitations in detecting protein products associated with NMD escape and resolving their underlying mechanisms, we combined a dual-color reporter system with fluorescence-activated cell sorting (FACS) and Western blotting to isolate cells exhibiting distinct NMD states and characterize the resulting protein outcomes. This reporter system uses a bidirectional promoter to simultaneously express wild-type and PTC-containing TPI reporters within the same cell, enabling systematic investigation of how PTC position influences NMD efficiency and downstream translational outcomes. This integrated approach enabled us to directly link PTC position to both mRNA fate and protein output following NMD escape, providing insight into potential mechanisms underlying NMD escape through distinct translational outcomes.

## Results

### A bidirectional reporter enables measurement of NMD, Premature termination, readthrough, and reinitiation downstream of PTC

To investigate how PTCs are resolved in human cells, we engineered a bidirectional dual reporter system that simultaneously quantifies mRNA decay and multiple translational outcomes, including premature termination, readthrough, and reinitiation, within a single reporter system. This system utilizes a Ponasterone A (PonA)-inducible bidirectional promoter to express two independent transcriptional units in opposite orientations from the same plasmid (Fig. 1A) ^37^. This inducible setup, combined with the bidirectional design, minimizes variability arising from transcription and transfection efficiency. In addition, the mCherry-TPI internal control provides a reference at both the RNA and protein levels, allowing for the normalization of GFP/mCherry ratios across cells.

**Figure 1.**
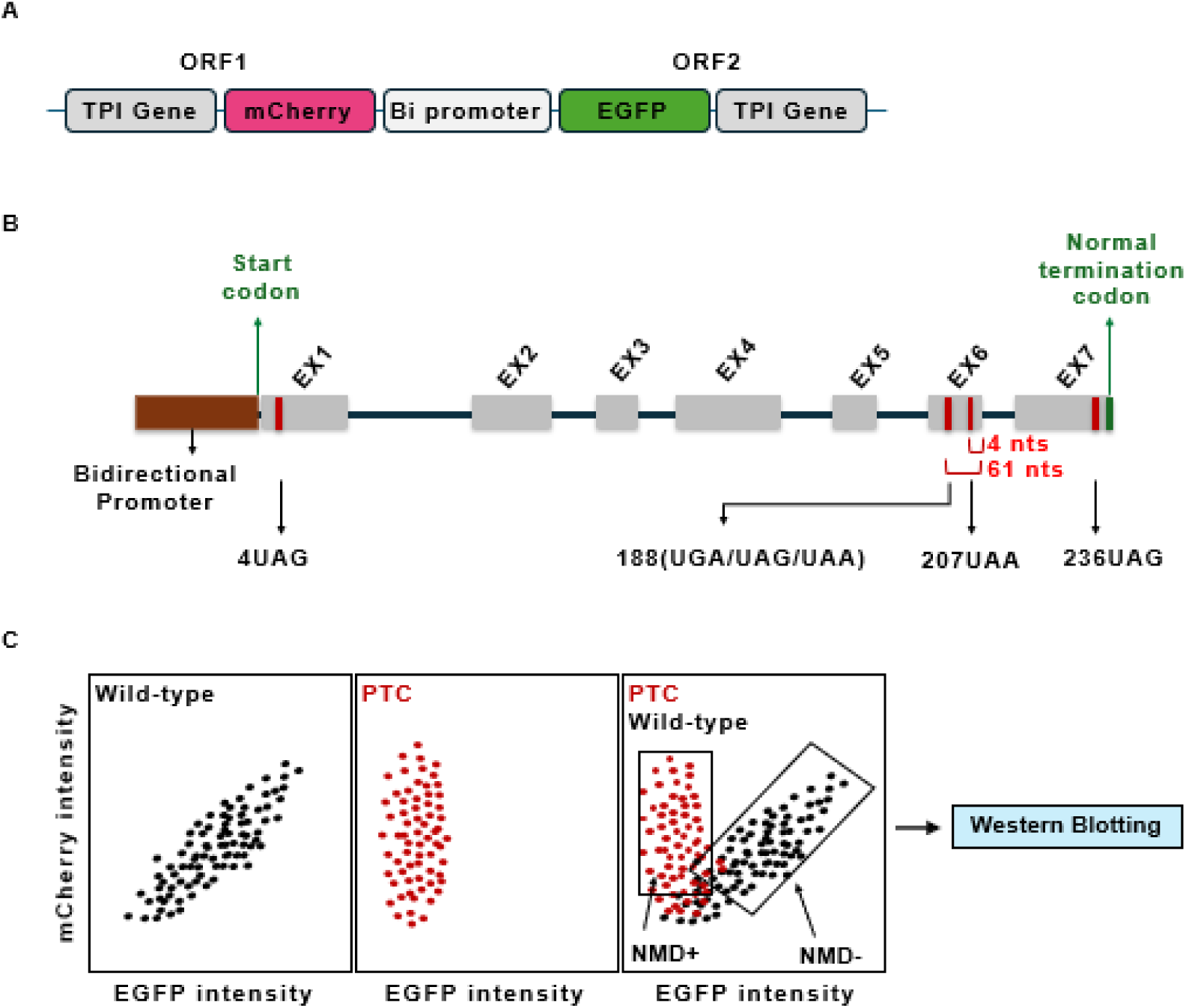
A bidirectional dual-color reporter system for single-cell NMD and protein outcome analysis. (A) A single promoter drives the simultaneous expression of an experimental reporter (GFP-TPI) and an internal control (mCherry-TPI) from a single plasmid. (B) Structural organization of the TPI reporter. The construct maintains the full exon–intron architecture and endogenous exon-exon junctions (EEJs) necessary for authentic NMD induction. (C) NMD efficiency is quantified at single-cell resolution by measuring GFP/mCherry ratios via flow cytometry. This allows for the isolation of cell populations with distinct NMD activities via FACS, followed by Western blot analysis to characterize the resulting protein outcomes.

Depending on the position of the GFP tag, distinct translation outcomes can be distinguished. The 5FP (GFP-TPI) reporter detects both premature termination and readthrough events, as GFP is produced in both cases, but does not capture translation reinitiation. In contrast, the 3FP (TPI-GFP) reporter detects readthrough and reinitiation events, while prematurely terminated products lacking the C-terminal tag are not detected.

We introduced premature termination codons (PTCs) at four patient-identified positions within the TPI coding region (codons 4, 188, 207, and 236) (Fig. 1B). Following transfection into HEK293T cells and induction with PonA, GFP and mCherry fluorescence intensities were measured by flow cytometry. Cells were then sorted into subpopulations exhibiting distinct NMD efficiencies (Fig. 1C), and the isolated populations were subsequently analyzed by Western blotting to characterize the resulting protein products.

### The type of stop codon does not affect NMD efficiency and its outcome at 188

We first examined whether the identity of the PTC influences NMD efficiency and protein outcomes in this reporter system. A PTC was introduced at codon 188 of the TPI gene (exon 6, ∼ 61 nucleotides upstream of the exon 6-7 junction), a location consistent with efficient NMD. To directly compare stop codon identity, all three stop codons (UGA, UAG, and UAA) were inserted at this site while maintaining identical flanking sequences (Fig. 2A).

**Figure 2.**
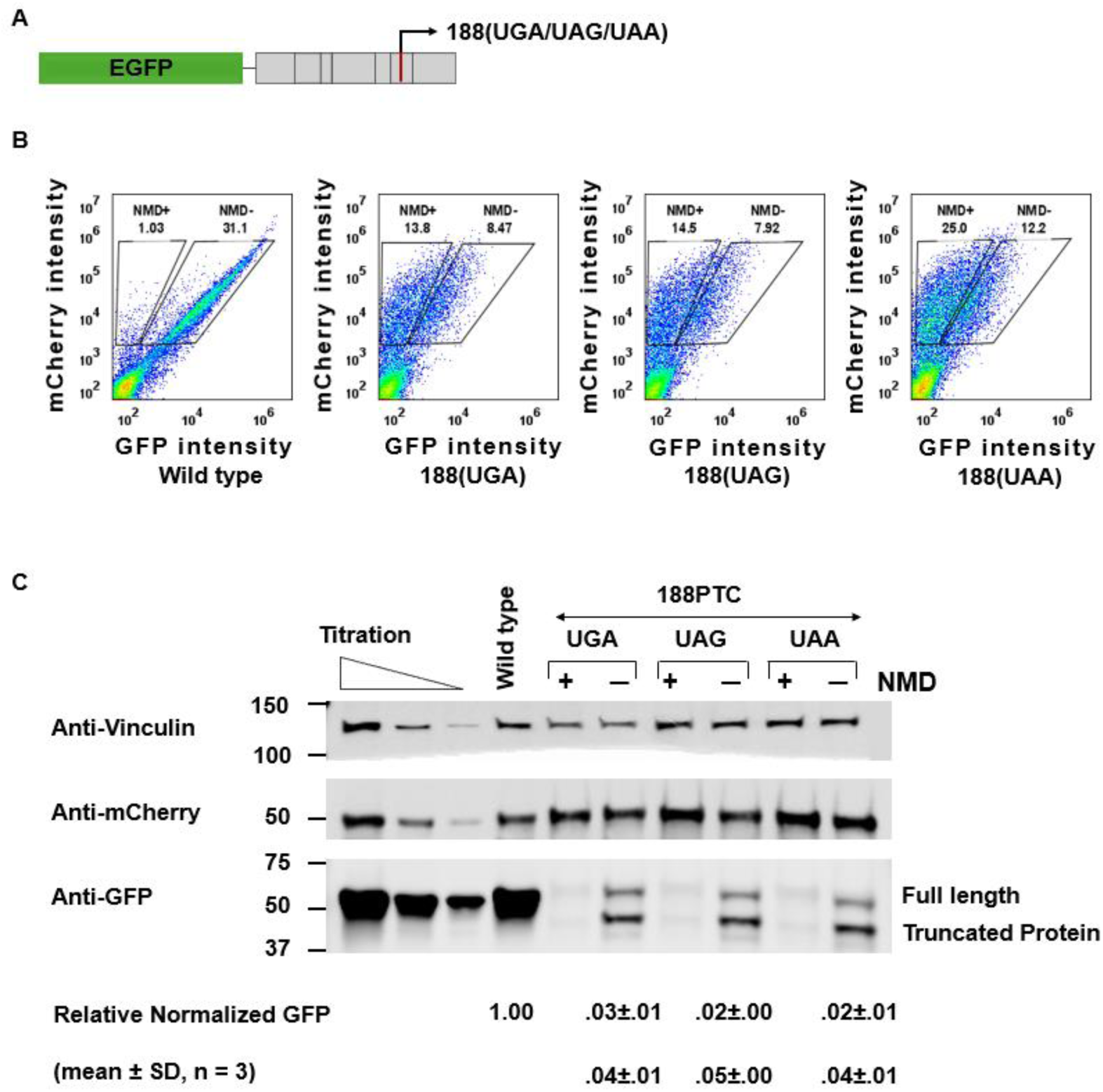
Stop codon identity has minimal impact on NMD efficiency and protein outcomes at position 188. (A) Schematic of GFP-TPI (5FP) reporter constructs. The GFP sequence is inserted upstream of the TPI coding sequence to monitor translational outputs. (B) Flow cytometry analysis of GFP-TPI (5FP) constructs. Flow cytometry of GFP-TPI (5FP) constructs reveals a significant reduction in GFP fluorescence across all PTC-containing mutants compared to the wild-type, indicating that NMD is triggered at this position. The decrease in GFP fluorescence reflects efficient mRNA degradation, a hallmark of NMD activity, as the GFP signal decreases in NMD positive (NMD+) cells. mCherry fluorescence from the internal control is unchanged and serves as an internal control for normalization. A reduced GFP/mCherry ratio corresponds to higher NMD efficiency, while higher ratios indicate NMD evasion, suggesting the presence of escaped transcripts. (C) Western blot analysis of protein isoforms. NMD-positive cells show a near-complete loss of full-length protein and the accumulation of truncated products. At codon 188, NMD is highly efficient regardless of the stop codon used (UAA, UAG, or UGA), indicating that stop codon identity has little influence on NMD strength or translational readthrough at this position. Relative protein levels were quantified by normalizing GFP band intensities to mCherry internal controls, with the wild-type ratio set to 1. Data are presented as mean ± SD from three independent experiments.

In the GFP-TPI (5FP) reporter, all three stop codons resulted in a marked reduction in GFP fluorescence, whereas mCherry expression derived from a wild-type, full-length control transcript remained unchanged (Fig. 2B). Normalization of GFP to mCherry showed a comparable decrease in the GFP/mCherry ratio across all stop codons, indicating similar NMD efficiency (Fig. 2C). Cells were further sorted by FACS into NMD-active (NMD⁺; low GFP) and NMD-escape (NMD⁻; high GFP) populations (Fig. 2B).

Western blot analysis using anti-GFP antibodies detected both full-length and truncated protein products, with the truncated species corresponding to termination at codon 188, whereas anti-mCherry antibodies detected only the full-length control protein (Fig. 2C). Protein profiles were comparable among the three stop codons, suggesting that both premature termination and readthrough occur with limited dependence on stop codon identity at this position. Quantification of bands revealed that approximately 2–3% of the protein corresponded to full-length products, consistent with readthrough events, whereas approximately 4–5% corresponded to truncated products arising from premature termination.

### Characterization of PTC Positions Predicted to Evade Start-Codon Proximal and EJC-dependent NMD

Next, we investigated PTC positions that are traditionally predicted to evade NMD through distinct mechanisms. These include codon 4, which is expected to bypass NMD due to its proximity to the start codon ^30^, as well as codons 207 and 236, which are evaluated in the context of the 50-55 nt rule governing EJC-dependent NMD ^14^ (Fig.3A). Notably, a UAA nonsense mutation at codon 207 in exon 6 is located 4 nucleotides upstream of the exon 6-7 junction, placing it near the boundary defined by the 50-55 nt rule. While this position is therefore predicted to be NMD-insensitive, its close proximity to the junction suggests that it lies at the margin between NMD-sensitive and -insensitive regions. In the GFP-TPI (5FP) reporter, the 207 UAA mutation showed a marked decrease in GFP fluorescence compared to the wild-type construct, as measured by flow cytometry, whereas mCherry fluorescence from the wild-type control transcript remained unchanged (Fig. 3B). These data suggest that NMD sensitivity is not defined by a strict positional cutoff but instead reflects a shift in the proportion of NMD-active and NMD-insensitive cells within the population. In contrast, most cells expressing UAG nonsense mutation at codon 236, located in the last exon where no downstream EJC is present, exhibiting high GFP expression, indicating minimal NMD activity. Together, these results indicate that NMD sensitivity transitions from a heterogeneous, non-binary state near the exon–exon junction to a uniformly NMD-insensitive state in the last exon.

**Figure 3.**
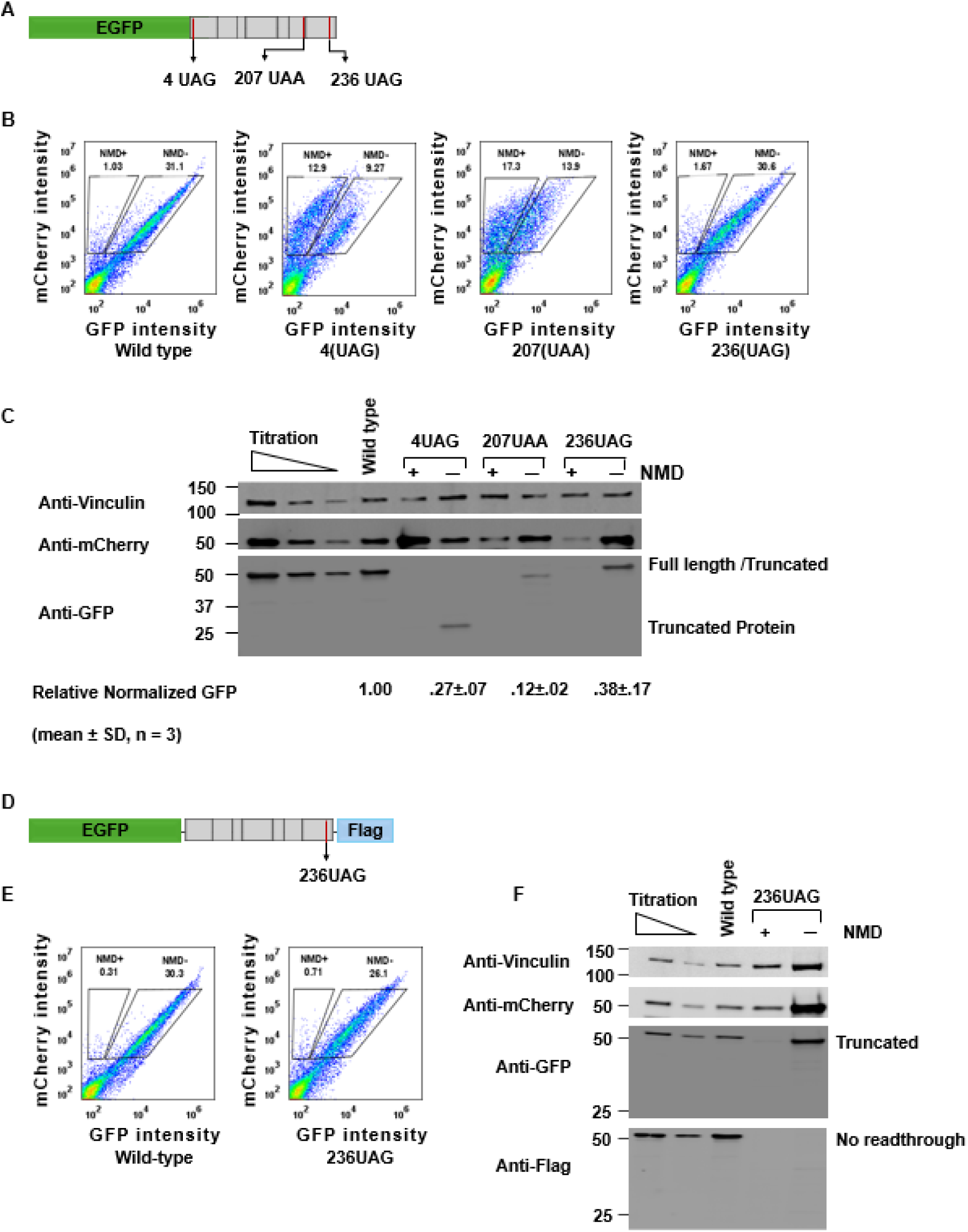
The 5FP reporter reveals protein outcomes from translational readthrough and premature termination at PTCs. (A) **Schematic of GFP-TPI (5FP) reporter constructs.** PTCs were introduced at defined positions within the TPI coding sequence to assess NMD efficiency and protein outcomes resulting from premature termination or translational readthrough. The GFP tag reports translational outputs, while mCherry serves as an internal control. (B) Flow cytometry analysis showing EGFP and mCherry fluorescence distributions for 5FP constructs containing various PTCs. Flow cytometry data show reduced GFP fluorescence in cells expressing PTCs compared to the wild-type (WT) control, indicating NMD activity at PTC-containing transcripts. The decrease in GFP fluorescence reflects mRNA degradation. mCherry fluorescence remains constant and is used for normalization. The GFP/mCherry ratio correlates with NMD efficiency, where lower ratios indicate higher NMD activity and higher ratios indicate NMD escape. (C) Western blot analysis of sorted cell populations with distinct NMD efficiencies. Cells were isolated based on distinct NMD efficiencies to resolve protein products under varying conditions. Western blots reveal truncated protein products generated by premature termination at the PTC, as well as full-length protein produced through translational readthrough. Truncated products correspond to termination at the PTC site, whereas full-length products indicate readthrough events. Relative protein levels were quantified by normalizing GFP band intensities to mCherry internal controls, with the wild-type ratio set to 1. Data are presented as mean ± SD from three independent experiments. (D) Schematic of C-terminally FLAG-tagged reporters. In GFP-TPI constructs, a FLAG tag was inserted at the 3′ end of the 5FP-236UAG construct to specifically track C-terminal extensions. (E) Flow cytometry analysis of FLAG-tagged GFP-TPI (5FP) 236UAG constructs. Fluorescence distributions for the 236UAG constructs used for downstream isolation. (F) Western blot analysis of sorted populations from FLAG-tagged constructs. Western blot analysis of sorted populations showing that specific protein products are derived from either translational readthrough (GFP-positive and FLAG-positive) or truncated products (GFP-positive and FLAG-negative) produced from premature termination.

To further examine how PTC position influences translation outcomes, including premature termination and readthrough, western blot analysis was performed using anti-GFP and anti-mCherry antibodies (Fig. 3C). This analysis revealed truncated protein products of the expected size corresponding to premature termination at codons 4 UAG and 207 UAA (Fig. 3C)

In contrast, for the codon 236 UAG mutation, the truncated product (52 kDa) could not be clearly distinguished from the full-length protein (53.38 kDa) due to its proximity to the normal stop codon. Quantitative analysis of the bands showed efficient production of a protein product in the NMD-escape cell population in the 236 UAG mutants (Fig. 3C).

To determine whether the protein produced in the 236 UAG mutant in the NMD-deficient cell population represents a truncated product resulting from premature termination or a full-length product arising from readthrough, GFP and FLAG tags were inserted at the N- and C-termini of the TPI reporter, respectively (Fig. 3D). In this system, readthrough products are expected to contain both FLAG and GFP, whereas premature termination products contain only the N-terminal GFP tag but lack FLAG. This design also enables evaluation with minimal alteration of the 3’sequence downstream of the 236 UAG codon due to the small size of the FLAG tag. FACS followed by Western blotting revealed the presence of a GFP-positive band but no corresponding FLAG signal, indicating that the protein is generated by premature termination rather than readthrough in the 236 UAG mutant (Fig. 3E, F).

### Translation Reinitiation Suppresses NMD and Promotes Readthrough

Proximity to the translation initiation codon reduces NMD efficiency, likely due to translation reinitiation downstream of the PTC ^35,36^. Although the 5FP reporter followed by Western blotting can distinguish translation outcomes such as premature termination and readthrough, it does not allow detection of N-terminally truncated products generated by translation reinitiation. To address translation outcomes from reinitiation, we generated a C-terminally tagged reporter by inserting GFP at the C terminus of the TPI gene (TPI-GFP; 3FP, Fig. 4A and Fig. S1A).

**Figure 4.**
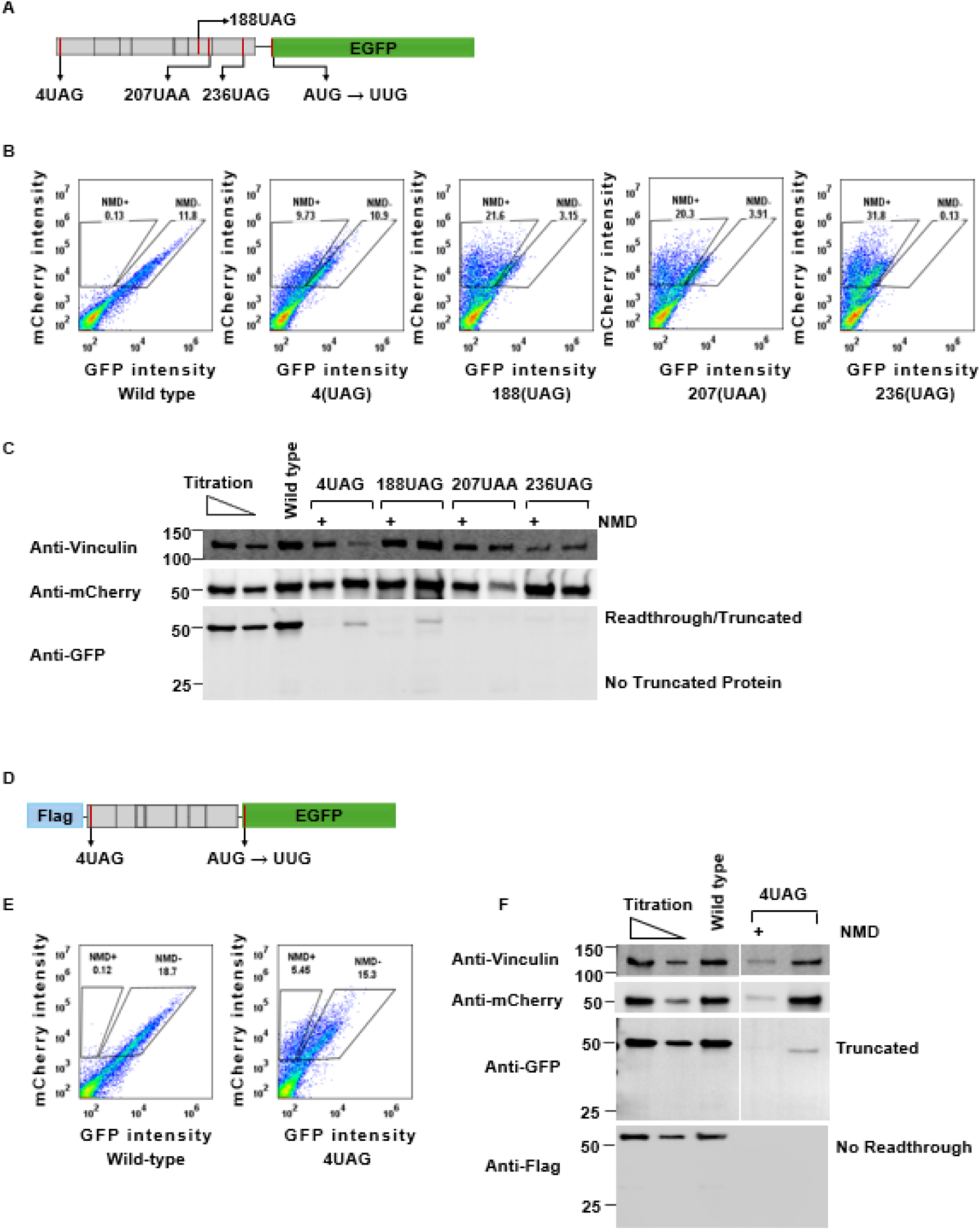
The 3FP reporter identifies protein outcomes from translational readthrough and downstream reinitiation. (A) Schematics of TPI-GFP (3FP) reporter constructs. PTCs were introduced at various positions within the TPI sequence, followed by a C-terminal GFP fusion to monitor translational outcomes. The GFP tag allows detection of reinitiation and readthrough products, while mCherry serves as an internal control for normalization. (B) Flow cytometry analysis of 3FP constructs containing various PTCs. Flow cytometry analysis shows EGFP and mCherry fluorescence distributions at the single-cell level. Cells with reduced NMD activity or increased translational readthrough/reinitiation exhibit higher GFP fluorescence, whereas NMD-active cells show lower GFP fluorescence. mCherry fluorescence remains constant across conditions and is used for normalization. The GFP/mCherry ratio correlates with NMD efficiency as well as translational readthrough or reinitiation outcomes. (C) Western blot analysis of sorted cell populations with distinct NMD efficiencies in 3FP reporters. Protein products from cell populations with distinct NMD efficiencies were analyzed to resolve isoforms generated by readthrough or reinitiation. (D) Schematic of N-terminally FLAG-tagged TPI-GFP constructs. A FLAG tag was inserted at the 5′ end of the 3FP-4UAG construct to specifically distinguish N-terminal extensions from internal initiation products. (E) Flow cytometry analysis of FLAG-tagged TPI-GFP (3FP) 4UAG constructs. Fluorescence distributions of the 3FP-4UAG constructs used for cell sorting and downstream analysis. (F) Western blot analysis of sorted populations from FLAG-tagged TPI-GFP 4UAG constructs. The results confirmed that the detected bands correspond to either full-length readthrough products (FLAG-positive) or N-terminally truncated reinitiation products (FLAG-negative).

When the downstream GFP retained its own initiation codon, a prominent band corresponding to GFP alone was unexpectedly detected in all mutants after cell sorting (Fig. S1B, C). This band was also observed in the wild-type construct, indicating that it occurs independently of the presence of a PTC. Although translation is generally thought to be initiated at the most upstream start codon via ribosomal scanning, these results suggest that initiation can also occur at the downstream GFP coding sequence in this context. While this observation deviates from the canonical scanning model and may reflect a context-dependent or non-canonical initiation event, mutation of the GFP start codon (AUG to UUG) completely abolished the corresponding band, confirming its origin (Fig. 4B, C).

The TPI coding sequence contains a natural AUG codon at amino acid position 14, as well as an additional downstream AUG at position 82. Among the constructs, only the 4 UAG mutant introduces additional in-frame initiation codons downstream, whereas the 188 UAG and other mutants do not. Western blot analysis showed that a TPI-GFP protein product was detected in the 4 UAG and 188 UAG mutants, but not in the 207 UAA and 236 UAG mutants. Because the protein produced by reinitiation in the 4 UAG mutant (51.89 kDa) is similar in size to the full-length TPI-GFP protein (53.38 kDa), we employed a tagging strategy to distinguish them. FLAG and GFP tags were inserted at the N- and C-termini of the TPI reporter, respectively (Fig. 4D). This design allows discrimination between reinitiation and readthrough products while minimizing perturbation of the native sequence context due to the small size of the FLAG tag. In this system, readthrough-derived products are expected to contain both the N-terminal FLAG and C-terminal GFP, whereas reinitiation-derived products contain only the C-terminal GFP tag. As shown in Fig. 4E and 4F, Western blotting after FACS revealed a GFP-positive band but no corresponding FLAG signal, indicating that the observed protein is generated by reinitiation. While both readthrough across the 4UAG codon and reinitiation at the AUG at position 14 were observed, no reinitiation products were detected from the AUG at position 82.

### NMD escape via translation reinitiation

Previous studies have shown that translation reinitiation downstream of a PTC can promote NMD escape in a distance-dependent manner ^30,32,35,36^. The results in Fig. 4D-F show that the AUG corresponding to amino acid position 14 (24 nt downstream of the PTC) supports protein production, whereas the AUG at position 82 (228 nt downstream of the PTC) does not. These findings suggest that reinitiation preferentially occurs at the first downstream AUG, while other in-frame AUG codons are either inactive or exhibit reduced efficiency in a context-dependent manner. Notably, the AUG at position 82 is located close to the exon-exon junction (7 nt upstream), raising the possibility that its proximity to the EJC may hinder efficient reinitiation.

Based on these observations, we next investigated how the position of downstream AUG codons influences reinitiation efficiency and the resulting protein products, as well as how the number of AUG codons affects NMD escape in the context of the 4 UAG mutation (Fig. 5A).

**Figure 5.**
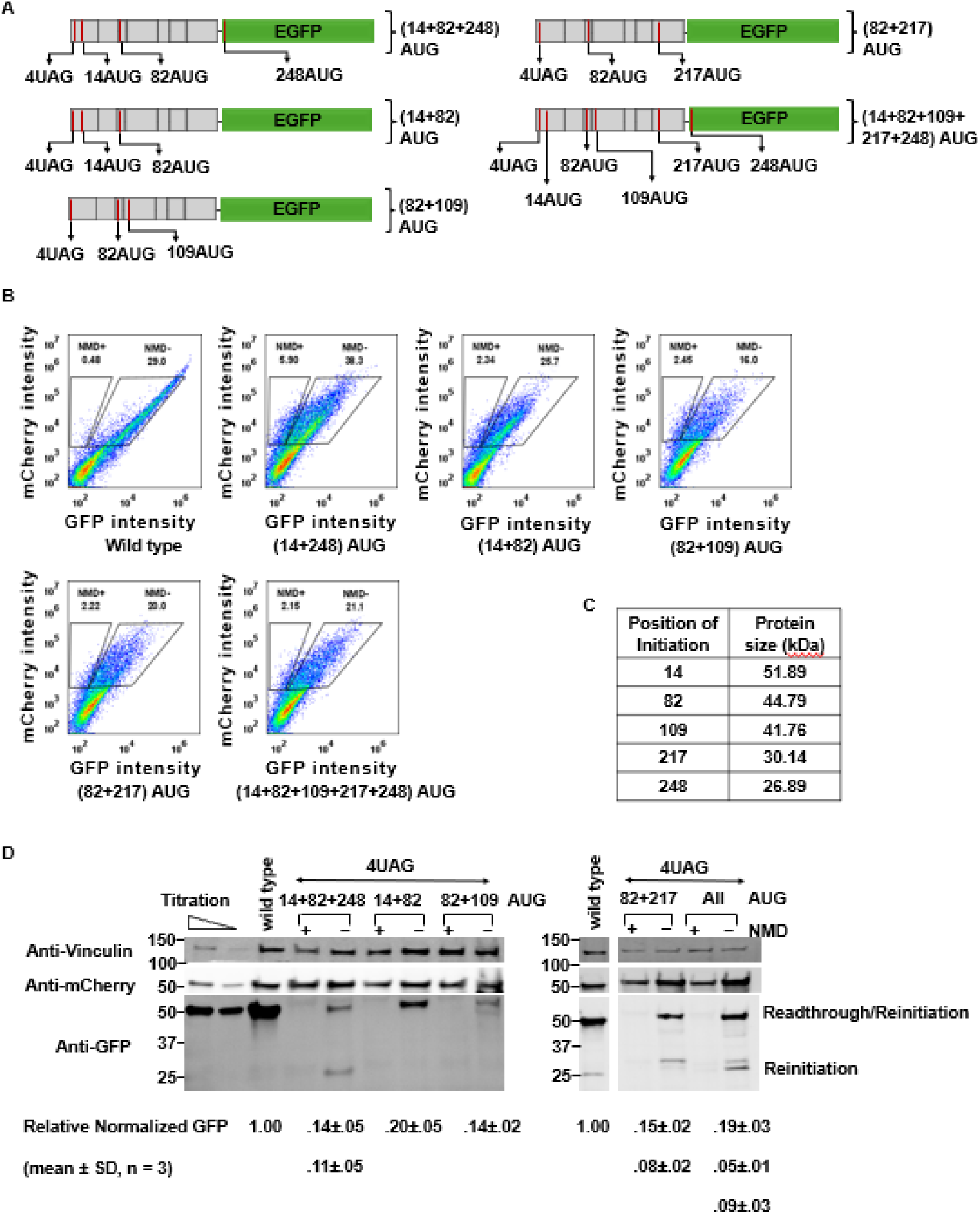
Influence of downstream AUG positions on NMD efficiency and protein outcomes via reinitiation. (A) Schematic of 4UAG PTC-containing TPI-GFP (3FP) constructs. A PTC (4UAG) was introduced at codon 4, and downstream AUG codons were inserted at defined positions within the TPI sequence to assess their effects on translation reinitiation and NMD outcomes. (B) Flow cytometry analysis of TPI-GFP (3FP) constructs. EGFP and mCherry fluorescence distributions are shown at single-cell resolution. The presence of downstream AUG codons (positions 14, 82, 109, 217, and 248) altered GFP fluorescence intensity, with higher GFP levels suggesting reduced NMD activity and/or increased translation reinitiation. These changes indicate that downstream initiation sites can modulate the translational output of PTC-containing transcripts. mCherry fluorescence remains constant across conditions and was used for normalization. (C) Predicted protein isoforms derived from downstream AUG usage. Theoretical molecular weights of protein products initiated from each downstream AUG codon are shown. Translation initiation at different AUG sites generates protein isoforms of distinct sizes depending on the position of reinitiation along the coding sequence. (D) Western blot analysis of reinitiation products. Immunoblotting of 3FP constructs reveals multiple protein species consistent with utilization of downstream AUG codons. GFP-positive bands correspond to full-length products, whereas lower molecular weight bands correspond to truncated proteins generated by reinitiation at downstream AUGs. Relative protein levels were quantified by normalizing GFP band intensities to mCherry internal controls, with the wild-type ratio set to 1. Data are presented as mean ± SD from three independent experiments.

To this end, we generated a series of constructs containing different combinations of downstream AUG codons in addition to the native sites at codons 14 and 82, including positions 109, 217, and 248 (corresponding to the native GFP start codon) (Fig. 5A).

FACS followed by Western blot analysis revealed that reinitiation at codon 82 is suppressed when an AUG is present at codon 14, regardless of the presence or absence of the AUG at codon 248 (Fig. 5B-D). In the absence of initiation at codon 248, which occurs independently of the PTC, protein production from reinitiation at codon 14 was increased compared to constructs retaining the AUG at codon 248, suggesting that limiting ribosome availability may segregate distinct protein outputs. In constructs containing an AUG at codon 109 or 217 but lacking the native AUG at codon 14, full-length readthrough products were detected. These products can be distinguished from reinitiation products based on their expected sizes (Fig. 5C), indicating that downstream AUG codons can promote readthrough even when located far from the PTC. In addition, protein production from translation initiation at codon 217 was also detected, suggesting that reinitiation does not always occur at the AUG closest to the PTC. When all downstream AUG codons were present, multiple protein products arising from both readthrough and reinitiation (e.g., from codons 217 and 248) were observed. These results indicate that post-termination ribosomes can scan across extended regions of the coding sequence and initiate translation at multiple downstream sites. Notably, these AUGs, except for the AUG at position 248, did not produce detectable protein in the wild-type reporter (Fig. S2), supporting that translation from these codons occurs primarily through PTC-dependent reinitiation rather than internal initiation.

### Reinitiation at downstream PTC attenuates NMD in a combinatorial and position-dependent manner

To determine how downstream PTC at 4 UAG influences NMD efficiency, we quantified reporter expression at the single-cell level using the EGFP/mCherry ratio. EGFP intensity was normalized to mCherry and further scaled to the median of the wild-type (WW) control within each experiment, followed by log₁₀ transformation.

We first compared NMD efficiency across constructs containing different combinations of AUG codons (Fig. 6A). Ranking based on median expression values revealed that constructs segregate into statistically distinct groups, indicating non-equivalent contributions of individual AUG configurations. Constructs containing all downstream AUG codons exhibited the highest normalized expression (ranking group “a”), consistent with the most efficient escape from NMD. In contrast, constructs containing AUGs at positions 82+109 or 82+217 showed the lowest expression levels (ranking group “c”), indicating limited escape from NMD. Notably, addition of the AUG at position 248 did not further increase expression compared to the (14+82) construct, as both conditions clustered within the same statistical group (ranking “b”), suggesting a minimal contribution of this site.

**Figure 6.**
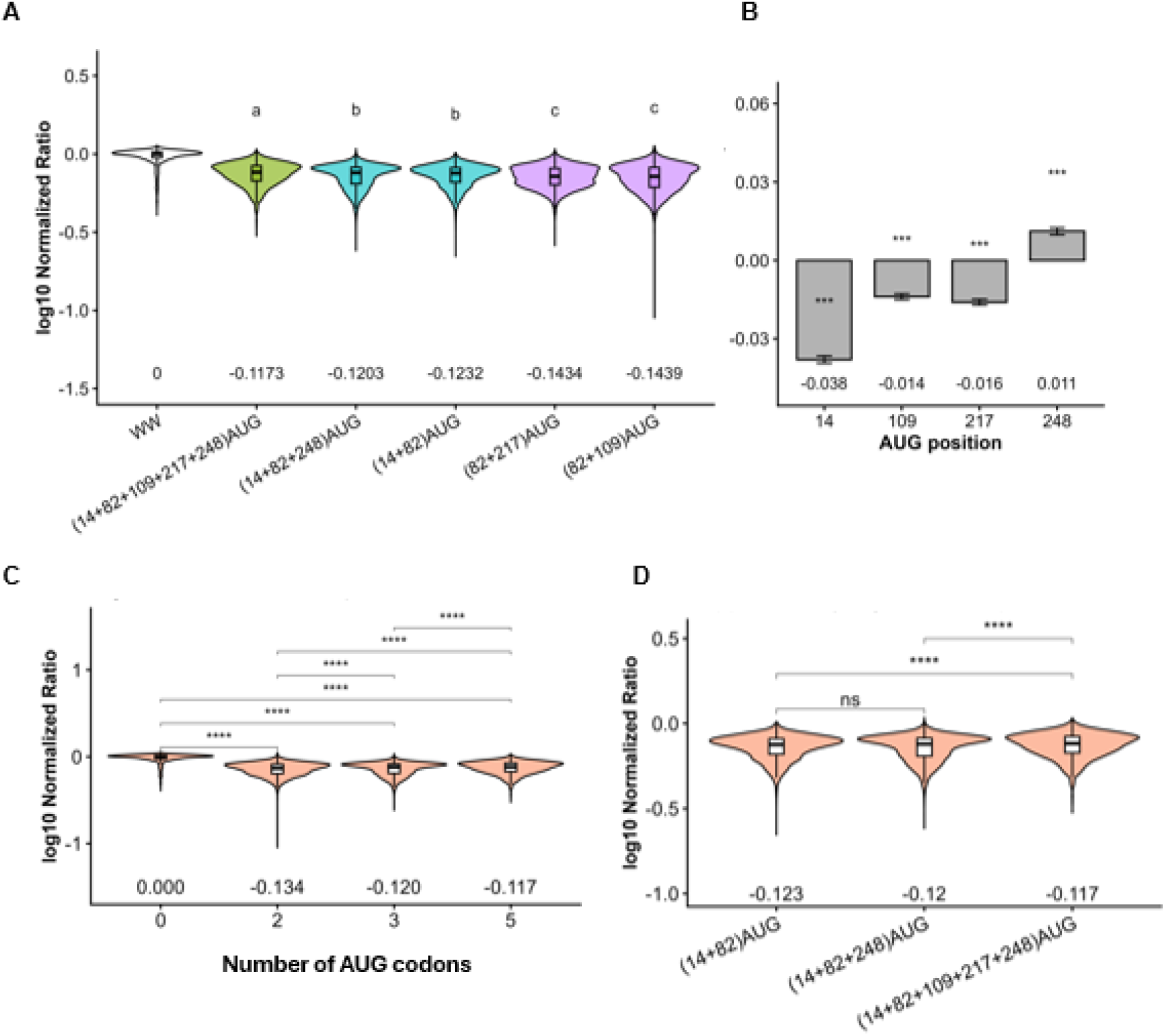
Combinatorial and positional effects of downstream AUG codons on NMD efficiency. (A) Ranked NMD efficiency across reporter constructs. For each cell, EGFP intensity was divided by mCherry intensity, and values were normalized to the median EGFP/mCherry ratio of the WW control within the same experiment, such that the WW median equals 1. The normalized ratios were log10-transformed for visualization. Distributions are shown as violin plots with overlaid boxplots. The median (log10 scale) for each condition is indicated below the plot. Statistical differences between groups were assessed using the Wilcoxon rank-sum test, with Benjamini-Hochberg correction for multiple comparisons. Groups not sharing the same letter are significantly different. Full results are provided in Supplementary Table 1. (B) Contribution of individual reinitiation AUG codons to NMD efficiency estimated by linear regression modeling. The dependent variable is the log10-transformed normalized EGFP/mCherry ratio. Each AUG position is treated as a binary variable (present = 1, absent = 0). Coefficient represents the effect size on log10 normalized expression, with error bars indicating standard errors. Statistical significance is indicated (*p < 0.05, **p < 0.01, ***p < 0.001). (C) Relationship between the number of upstream AUG codons (AUG count) and NMD efficiency. Normalization and transformation were performed as in (a). Violin plots show the distribution of log10 normalized ratios, with median values indicated. Sample size (n) is shown for each group. Statistical comparisons were performed using the Wilcoxon rank-sum test with multiple comparison corrections, and significance is indicated on the plot. (D) Stepwise addition of reinitiation AUG codons reveals additive and saturating effects on NMD efficiency. Normalization and log10 transformation were performed as described in (a). Median values are shown, and statistical comparisons between selected conditions were performed using the Wilcoxon rank-sum test. Adjusted p values are shown as significance levels, and full results are provided in Supplementary Table 1.

To quantify the contribution of individual AUG codons, we performed linear regression analysis using the presence or absence of each AUG as predictors (Fig. 6B). Because the AUG at position 82 was present in all constructs and appeared functionally silent (Fig. 5), it was excluded from the model. This analysis revealed a strong effect of the AUG at position 14, whereas AUGs at positions 109 and 217 contributed more modestly to NMD escape, indicating position-dependent differences in reinitiation efficiency.

We next examined whether the number of AUG codons correlates with NMD attenuation. Increasing AUG count was associated with a progressive increase in normalized expression (Fig. 6C), suggesting that multiple reinitiation sites act cumulatively to suppress NMD. However, deviations from a strictly linear trend indicate that these effects are not purely additive.

Stepwise comparison of constructs with incremental addition of AUG codons showed that the AUG at position 14 exerted the strongest effect on reporter expression, whereas AUGs at positions 109 and 217 contributed more modestly, and the AUG at position 248 had minimal impact (Fig. 6D). While the addition of individual sites resulted in only limited increases in expression, their combined presence produced a statistically significant enhancement, indicating cooperative contributions to NMD escape. Notably, the incremental gain in expression diminished with increasing AUG number, suggesting a saturating effect. Together, these results demonstrate that downstream AUG codons modulate NMD efficiency through position-dependent and cooperative mechanisms, with saturation limiting the overall extent of NMD attenuation.

### Positional NMD variability landscape

The strength of this approach is single-cell resolution, which enables us to investigate the cell-to-cell variability of NMD. NMD activity at single-cell resolution was inferred from protein output, quantified as the normalized EGFP/mCherry fluorescence ratios in individual cells expressing 5FP or 3FP reporters under various nonsense mutation conditions, including multiple PTC sites and the wild-type control (WW). These ratios reflect protein production outcomes following NMD escape rather than NMD efficiency per se.

Violin plot analysis revealed that all nonsense mutation constructs exhibited reduced normalized ratios compared to WW, consistent with NMD-mediated suppression of protein output. At the same time, substantial heterogeneity across individual cells was observed (Fig. 7A-B), and the fraction of escape cells was quantified for each condition (Fig. 7C-D). Using a WW-based threshold defined as the 5th percentile of normalized ratios in control cells, individual cells were classified as “Escape” or “Non-escape” (red lines in Fig. 7A-B; red bars in Fig. 7C-D).

**Figure 7.**
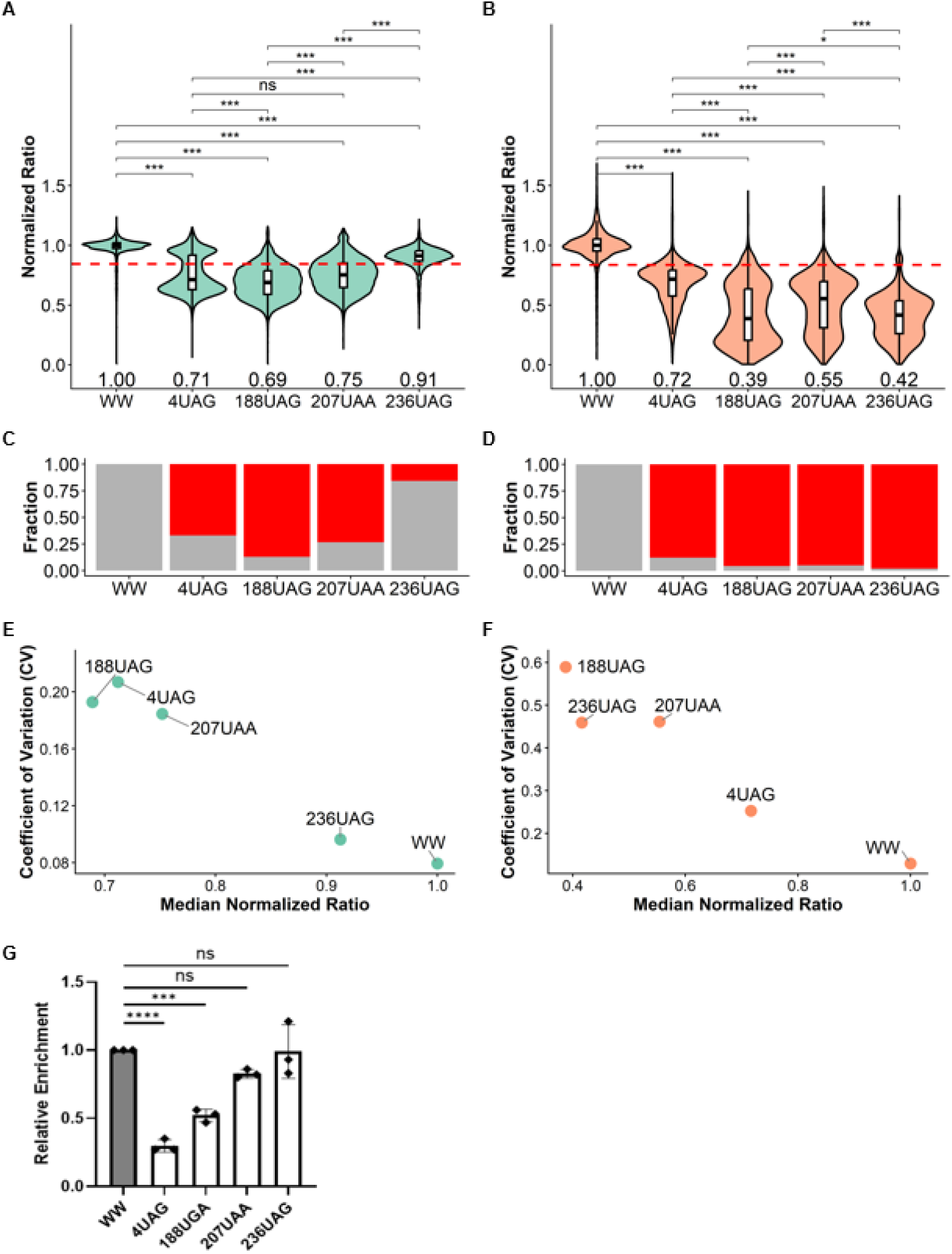
Single-cell variability and stochastic escape from NMD. (A-B) Distribution of NMD reporter expression across nonsense mutation conditions. EGFP fluorescence was normalized to mCherry at the single-cell level and further normalized to the median EGFP/mCherry ratio of the wild-type (WW) control within each experiment. Violin plots with overlaid boxplots show the distribution of normalized ratios for 5FP (A) and 3FP (B) reporters. Median values are indicated. Each condition corresponds to a distinct premature termination codon (PTC) position. Sample size (n) represents the number of single cells analyzed per condition. Statistical differences were assessed using pairwise Wilcoxon rank-sum tests with Benjamini-Hochberg correction. Adjusted p values are shown as significance levels, and full results are provided in Supplementary Table 1. A dashed line indicates the WW-derived escape threshold (5th percentile). (C-D). Fraction of NMD escape at the single-cell level. Cells were classified as “escape” or “non-escape” based on whether their normalized ratio exceeded the WW-derived 5th percentile threshold. Bar plots show the fraction of escape (red) and non-escape (gray) cells for each condition in 5FP (C) and 3FP (D) reporters. (E-F). Relationship between NMD efficiency and cell-to-cell variability. Scatter plots show the median normalized ratio (NMD efficiency) versus coefficient of variation (CV) for each condition in 5FP (E) and 3FP (F) reporters. Each point represents a condition. These plots highlight differences in both the magnitude and heterogeneity of NMD across PTC positions. (G) Quantitative RT-PCR analysis of reporter transcript abundance. Relative gene expression in wild-type and PTC-containing constructs was measured, with WT serving as the reference control. Data are presented as mean ± SEM from three biological replicates (n=3). Statistical significance was determined by one-way ANOVA followed by Dunnett’s multiple comparison test. Statistical significance was determined relative to WT. ***p < 0.001, ****p < 0.0001; ns, not significant.

In the 5FP reporter, a substantial fraction of cells escaped NMD in the 236UAG construct, consistent with the 55-nt rule (Fig. 7A, C). In contrast, the 188UAG and 207UAA constructs exhibited broad unimodal distributions, whereas the 4UAG construct showed a clear bimodal distribution. A unimodal distribution suggests continuous variation in protein output, likely driven by quantitative differences such as translation efficiency or NMD factor abundance. In contrast, the bimodal distribution observed for 4UAG indicates the presence of discrete cellular states, consistent with switch-like regulation of NMD escape.

In the 3FP reporter, where GFP expression depends on translation of the downstream region, most mutation constructs exhibited reduced normalized ratios, likely reflecting effective 3′UTR extension caused by GFP insertion (Fig. 7B, D). Notably, the 188UAG, 207UAA, and 236UAG constructs lack in-frame downstream reinitiation codons. Therefore, GFP production in these contexts primarily reflects translational readthrough rather than reinitiation. In contrast, the 4UAG construct likely produces GFP via a combination of reinitiation and readthrough mechanisms.

As a result, in the 3FP system, protein output becomes partially uncoupled from NMD escape, as cells that evade NMD do not necessarily produce detectable GFP.

Cumulative distribution functions (CDFs), which show the fraction of cells below a given protein level (Fig. S3), and Z-score density plots (Fig. S4), which normalize values relative to the WW mean, further illustrated the distributions of single-cell responses. Compared to the 5FP reporter, the 3FP system exhibited broader CDF profiles across multiple conditions, indicating increased variability in protein output. Z-score density analysis revealed distinct modes of perturbation between reporters. In the 5FP system, bimodal distributions were observed in specific conditions, reflecting discrete escape states. In contrast, the 3FP system remained largely unimodal across conditions, with increased spread rather than clear subpopulations.

In the 3FP context, variability arises from continuous fluctuations in protein output rather than discrete switching between NMD-active and NMD-escaped states. The discrepancy in distribution patterns between 5FP and 3FP reporters is attributable to differences in how protein output is generated. In particular, NMD escape coupled with translational reinitiation can produce N-terminally truncated proteins that are GFP-negative in the 5FP system but GFP-positive in the 3FP system, highlighting how reporter design influences the detection of NMD escape events.

To directly examine the relationship between protein output and variability, we plotted the median normalized ratio against the coefficient of variation (CV) for each condition (Fig. 7E-F). In the 5FP reporter, higher median ratios were associated with increased variability, indicating that NMD escape enhances cell-to-cell heterogeneity. In contrast, in the 3FP reporter, the absence of downstream reinitiation codons uncouples NMD escape from GFP production, such that cells that escape NMD do not necessarily exhibit increased fluorescence, resulting in compression of the dynamic range and an apparent increase in variability.

To validate our single-cell findings using a conventional approach, we measured 5FP reporter mRNA levels by RT-qPCR (Fig. 7G). Overall, the RT-qPCR results were consistent with the GFP/mCherry ratios in 5FP reporters, except for the 4UAG construct. Although 4UAG was expected to escape NMD because of its proximity to the start codon, RT-qPCR showed significantly reduced mRNA levels in the 5FP reporter. This is likely because insertion of GFP upstream of TPI increases the distance between the start codon and the 4UAG PTC. As a result, 4UAG no longer behaves as a start-proximal PTC, but rather as an internal PTC that is more sensitive to NMD, consistent with our previous results (Fig. 3).

To quantitatively evaluate these trends, we summarized key statistical parameters for each condition, including median normalized ratio, variance, and coefficient of variation (CV) (Table 1). Pairwise comparisons using Wilcoxon rank-sum tests confirmed significant differences in protein output between WW and nonsense mutation constructs, as well as between specific mutation sites. These results underscore that protein output following NMD escape is both construct- and site-dependent, and that single-cell analysis reveals substantial heterogeneity in these outcomes.

**Table 1.**
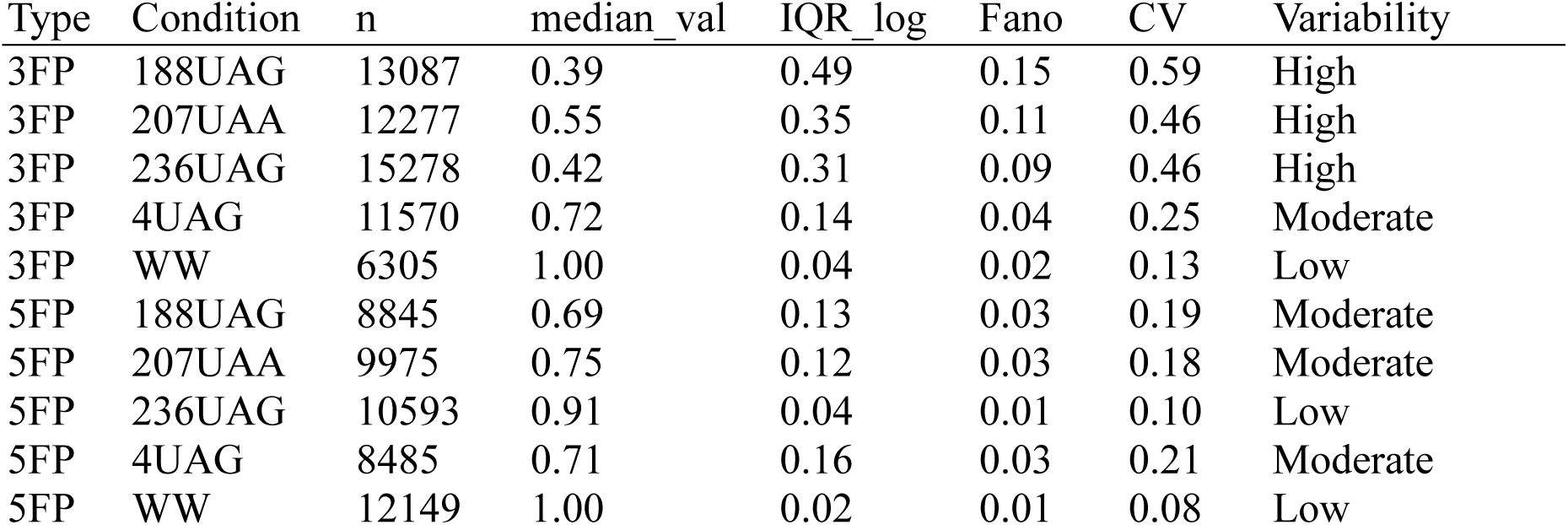
Variability metrics of NMD reporter constructs.

Together, these results demonstrate that NMD escape is not only associated with changes in overall protein output but also with shifts in the fraction of escape-prone cells and increased variability across the population. These findings highlight the inherently heterogeneous and context-dependent nature of NMD regulation at the single-cell level.

## DISCUSSION

In this study, we utilized a bidirectional dual-reporter system based on the TPI NMD reporter that enables simultaneous detection of NMD activity and protein output from NMD-escaping transcripts, including products generated by premature termination, translational readthrough, and reinitiation, at single-cell resolution. While determinants influencing NMD susceptibility, including EJC positioning, 3‘UTR length, and proximity to the start codon, the downstream consequences of these escape events have remained unclear. In particular, whether NMD-escaping transcripts are translated and what types of protein products they generate has not been well defined, in part due to the limited sensitivity of conventional approaches. Because NMD escape occurs in only a subset of cells, these events are difficult to resolve at the population level.

This bidirectional design overcomes these limitations by integrating both experimental and internal control transcripts within a single construct while preserving the native exon–intron structure. This design enables robust normalization, ensures comparable splicing and cellular context, and allows simultaneous quantification of mRNA stability together with multiple protein outputs, including full-length, truncated, and reinitiation-derived products. By directly linking transcript fate to protein output, this system provides a framework to examine how PTC-containing transcripts are interpreted by translation and mRNA surveillance machinery.

Our results reveal that the consequences of PTCs are more diverse than previously appreciated. PTC position, EJC context, and downstream sequence features collectively determine multiple outcomes, including mRNA decay, truncated protein production, translational readthrough, and reinitiation.

This positional dependence was particularly evident in translation reinitiation. Previous studies have reported that downstream initiation codons located approximately 20–30 nucleotides from a PTC can promote translation reinitiation and attenuate NMD ^30,32,35,36^. Consistent with these observations, the downstream AUG at position 14 exerted the strongest effect on NMD attenuation. By contrast, more distal AUGs (positions 109, 217, and 248) showed progressively weaker effects on NMD attenuation despite contributing to protein production. Systematic comparison of multiple initiation sites further demonstrated that reinitiation efficiency is strongly constrained by AUG position. While individual AUGs had limited effects, their combined presence increased protein expression in an additive manner, although this effect showed signs of saturation with increasing AUG number. These findings indicate that proximal downstream initiation sites near the PTC (∼20 nucleotides) are most effective in promoting NMD escape, whereas distal initiation sites primarily modulate translational output without equivalently affecting NMD efficiency.

Together, our findings support a model in which NMD escape and translational outcomes are governed by a shared termination environment rather than a single linear pathway. Translation downstream of a PTC may alter local mRNP architecture by changing ribosome occupancy and the spatial relationship between the termination codon and PABPC1. In parallel, competition between release factors and near-cognate tRNA incorporation, together with the availability of translation initiation and termination factors, may determine whether termination proceeds canonically or is diverted toward readthrough or reinitiation. In addition, translation may modulate accessibility of NMD factors such as UPF1 and EJCs, while temporal competition between pioneer-like and steady-state translation further shapes NMD outcomes.

At the population level, these combinatorial effects translate into heterogeneous cellular responses. Single-cell analysis revealed that NMD produces a spectrum of outcomes, ranging from efficient decay to escape. In the 5FP reporter, certain constructs exhibited bimodal distributions consistent with discrete NMD-active and NMD-escaped states, whereas others showed unimodal distributions reflecting continuous variation in protein output. These results indicate that NMD can display both switch-like and graded behaviors depending on transcript context.

The 3FP reporter further demonstrates that variability in protein output is shaped not only by NMD efficiency but also by how protein production is mechanistically coupled to NMD escape. In this system, the absence of downstream reinitiation codons uncouples NMD escape from GFP production, resulting in broader distributions without distinct subpopulations. This suggests that apparent cellular variability can arise from translational constraints rather than differences in NMD state per se.

Importantly, differences between the 5FP and 3FP reporters underscore how reporter design influences the interpretation of NMD activity. Reinitiation can generate N-terminally truncated proteins that are differentially detected depending on reporter configuration, leading to apparent differences in variability and escape frequency. These observations emphasize the need to jointly consider RNA decay and translational outputs when assessing NMD.

More broadly, our results suggest that NMD should not be viewed as a strictly binary quality-control pathway, but rather as a probabilistic process shaped by transcript architecture and translational context. The combination of position-dependent reinitiation and stochastic cellular variation generates diverse protein expression outcomes from identical transcripts, which may contribute to phenotypic heterogeneity in genetic disorders caused by nonsense mutations.

Future studies applying this system across different genes and cell types will help define how PTC outcomes are modulated by sequence and cellular context. Identifying factors that influence the relative contributions of mRNA decay, truncation, readthrough, and reinitiation will be critical for understanding gene expression control and may inform therapeutic strategies aimed at restoring functional protein expression.

## Material and Methods

### Plasmid construction

All plasmid used in this experiment were generated by standard molecular cloning procedures. The human triosephosphate isomerase (TPI) gene was amplified from the plasmid (pcDNA5/FRT/TO-TPI_WT-xrRNA-4H, TPI1 Human, addgene-plasmid, #108377) by polymerase chain reaction (PCR) using gene specific primers containing appropriate restriction enzyme recognition sites and cloned into pFRT-PonA-BI-TPI-GFP-WT-mCherry-WT either downstream of GFP (GFP-TPI, 5FP construct) or upstream of GFP (TPI-GFP, 3FP construct) using standard restriction enzyme based cloning. PCR products and vectors (Ponasterone A inducible bidirectional reporter plasmid, Sato and singer., Nat comm) were digested with the indicated enzymes, purified, and assembled using 10 μl of 2X Gibson Assembly (NEBuilder HiFi DNA Assembly Master Mix, NEB #E2621S, New England Biolabs), in a total of 20 μl at 50°C for 60 minutes or Ligation-Convenience Kit (#319-05961, Nippon Gene, Tokyo, Japan). Dual reporter plasmids were constructed by inserting the EGFP-TPI and mCherry-TPI expression cassettes in opposite orientations under the control of the PonA inducible bidirectional promoter ^13^. Nonsense mutation constructs were generated by overlap extension PCR using primers containing the desired nucleotide substitutions. For reinitiation analysis, downstream AUG codons were introduced to evaluate translation reinitiation. Transformation was performed using TOP10 Chemically Competent E. coli (Thermo Fisher Scientific Inc.), Plasmid DNA was isolated from bacterial cultures using the FastGene Plasmid Mini Kit (FG-90502; Nippon Genetics, Tokyo, Japan), screened by restriction digestion, and confirmed by Sanger sequencing across the entire insert and cloning junctions.

### Cell culture and induction

#### Cell line and maintenance

HEK293T PonA ^37^ cells were used for all experiments. Cells were cultured in Dulbecco’s Modified Eagle Medium (DMEM) (4.5 g/l glucose, Nacalai Tesque Co., Ltd.) supplemented with 10% fetal bovine serum (FBS) SA (Cat. No. 175012-500ML; Nichirei Biosciences, Tokyo, Japan) and 1% penicillin streptomycin (Cat. No. 06168-34, Nacalai Tesque Co., Ltd.) at 37° C in a humidified incubator with 5% carbon dioxide. Cells were routinely passaged at 70-90% confluence.

#### Ponasterone A (PonA) inducible expression

All reporter constructs were expressed using a bidirectional promoter responsive to Ponasterone A. To induce protein expression, after 4-6 hours post transfection, cells were treated with 5 μM Ponasterone A (Cat. No. 16386; Cayman Chemical) for 24 hours. Induction was allowed to proceed for 24-48 hours prior to analysis.

#### Transfection

Reporter constructs were transiently transfected into HEK293T PonA cells using Lipofectamine 2000/3000 Transfection Reagent (Cat. No. 13778100; Invitrogen/Thermo Fisher Scientific) or PEI MAX-Transfection GradeLinear Polyethylenimine Hydrochloride (Cat. No. 24765-100; Polysciences, Inc.) according to the manufacturer’s instructions. For transfection, cells were seeded at ∼60-70% confluence and transfected with plasmid DNA. Transfection complexes were prepared in Opti-MEM™ I Reduced Serum Medium (Cat. No. 31985062; Gibco/Thermo Fisher Scientific), incubated at room temperature for 15-20 min to form complexes, and added dropwise to cells. All constructs (wild-type and mutant-type) were transfected in parallel using identical conditions to ensure consistency. For experiments requiring higher cell numbers (FACS sorting or Western blot), cells were transfected in 10 cm dishes with 12-15 ug DNA while maintaining the same DNA to reagent ratio.

#### Fluorescence-activated cell sorting (FACS)

At 24–48 h after ponasterone A (PonA) induction, cells were washed with Dulbecco’s phosphate-buffered saline without calcium and magnesium (D-PBS(–); Nacalai Tesque) and dissociated using 2.5 g/L trypsin and 1 mmol/L EDTA solution with phenol red (Nacalai Tesque) for 5 min at 37°C. Cells were resuspended in ice-cold PBS supplemented with 2% fetal bovine serum (FBS) to maintain viability. Cell suspensions were passed through a cell strainer to remove aggregates and ensure single-cell measurements.

Flow cytometry and cell sorting were performed using a SH800 Cell Sorter (Sony Biotechnology). EGFP and mCherry fluorescence were detected using 488 nm and 561 nm excitation lasers, respectively, with consistent gating applied across all experiments. A minimum of 50,000 cells per sample was collected for quantitative analysis. For biochemical analysis, cells were sorted into two populations based on EGFP intensity under identical gating conditions: NMD-positive (NMD+) cells, defined by low EGFP signal (indicative of efficient mRNA decay), and NMD-negative (NMD–) cells, defined by high EGFP signal (indicative of stabilized transcripts). For each population, 5 × 10⁴ to 1 × 10⁵ cells were collected directly into ice-cold PBS containing 2% FBS and immediately processed for downstream protein extraction. All experiments were performed with at least three independent biological replicates, each derived from separate transfections and inductions.

#### Flow cytometry data processing and normalization

Flow cytometry data were exported as CSV files and analyzed in R (v4.x). Single-cell fluorescence intensities for EGFP and mCherry were extracted. Cells were filtered to exclude low-signal and saturated events (EGFP ≤ 1 or ≥ 9.5 × 10⁵; mCherry ≤ 1 or ≥ 9.5 × 10⁵).

To account for background autofluorescence, the median EGFP signal from non-transfected control samples was calculated. Reporter expression was quantified as the ratio of EGFP to mCherry fluorescence for each cell. These ratios were normalized to the median EGFP/mCherry ratio of the wild-type (WW) control within the same experiment, thereby setting the WW median to 1.

For analyses requiring stabilization of variance, normalized ratios were log₁₀-transformed.

#### Analysis of reinitiation-dependent effects on NMD (Figure 6)

Constructs containing different combinations of upstream AUG codons were analyzed to assess the impact of translation reinitiation on NMD efficiency.

Normalized EGFP/mCherry ratios were log₁₀-transformed and visualized using violin plots with overlaid boxplots.

#### Ranking analysis

Constructs were ranked based on the median log₁₀-transformed normalized ratio. Pairwise comparisons between all conditions were performed using the Wilcoxon rank-sum test with Benjamini–Hochberg correction for multiple testing.

Adjusted p values were used to generate a compact letter display, in which constructs sharing the same letter are not significantly different. Letters were reassigned according to descending median values, such that the highest-expression group was labeled “a,” followed by “b,” “c,” and so on.

#### Contribution of individual AUG codons

To quantify the contribution of each reinitiation AUG codon, a linear regression model was fitted using the presence or absence of individual AUG sites as independent variables and the log₁₀-transformed normalized ratio as the dependent variable. Regression coefficients represent effect sizes, and standard errors were calculated for each estimate.

#### Cumulative and stepwise analyses

To evaluate cumulative effects, constructs were grouped by the total number of AUG codons and compared using the Wilcoxon rank-sum test with multiple testing correction.

To assess additivity and potential saturation, selected constructs with stepwise addition of AUG codons were compared using the same statistical framework.

#### Analysis of single-cell variability and NMD escape

To characterize cell-to-cell heterogeneity in NMD, analyses were performed on normalized (non-log-transformed) EGFP/mCherry ratios.

#### Distribution analysis

Cumulative distribution functions (CDFs) were computed for each condition to evaluate differences in distribution shape and tail behavior.

#### Variability metrics

Cell-to-cell variability was quantified using the coefficient of variation (CV; standard deviation divided by mean) and Fano factor (variance divided by mean).

#### Definition of NMD escape

An empirical threshold for NMD escape was defined using the wild-type (WW) distribution. Specifically, the 5th percentile of the WW normalized ratio was used as a cutoff. Cells with normalized ratios above this threshold were classified as “escape,” whereas those below were classified as “non-escape.”

#### Escape fraction analysis

The fraction of escape cells was calculated for each condition and visualized as proportional bar plots.

#### Z-score normalization

For distribution-based comparisons, log₁₀-transformed normalized ratios were converted to Z-scores across the dataset, and density plots were used to compare shifts in population distributions.

#### Statistical analysis (Figure 6 and 7)

All statistical analyses were performed in R. Differences between groups were assessed using the Wilcoxon rank-sum test. P values were adjusted for multiple comparisons using the Benjamini– Hochberg method.

Statistical significance is reported as adjusted p values (*p < 0.05, **p < 0.01, ***p < 0.001). Sample size (n) represents the number of single cells analyzed per condition. Median values are reported unless otherwise specified.

#### Data visualization (Figure 6 and 7)

Data were visualized using ggplot2. Violin plots with overlaid boxplots were used to display distributions. Statistical significance was annotated based on adjusted p values. Additional visualizations included cumulative distribution plots, density plots, and proportional bar plots.

#### Cell lysis and Western blotting

After cell sorting, cells were washed twice with cold PBS and lysed on ice in Pierce IP Lysis Buffer (Thermo Fisher Scientific) (25 mM Tris-HCl, pH 7.4; 150 mM NaCl; 1% NP-40; 1 mM EDTA; 5% glycerol) supplemented with protease inhibitor cocktail (cOmplete Mini, Roche). Lysates were incubated on ice for 10 min and clarified by centrifugation at 13,000 × g for 10 min at 4°C, and the supernatant containing soluble proteins was collected. Protein samples were mixed with 4× Laemmli Sample Buffer (Bio-Rad) and stored at −30°C until use. Equal amounts of protein were boiled at 95°C for 5 min and separated by SDS–PAGE, followed by transfer onto a 0.2 µm Immobilon-PSQ PVDF membrane (Merck Millipore). Membranes were blocked for 30 min at room temperature in blocking buffer (EveryBlot Blocking Buffer, Bio-Rad) and incubated overnight at 4°C with primary antibodies against GFP (to detect full-length, truncated, and reinitiation-derived products), mCherry (internal control), FLAG (to detect tagged full-length and truncated proteins), and vinculin (loading control). After washing with TBS-T, membranes were incubated with IRDye-conjugated secondary antibodies (LI-COR) for 1 h at room temperature, and protein bands were visualized using an Odyssey F Imaging System (LI-COR Biosciences) with near-infrared detection at 700 and 800 nm.

#### Band analysis

Immunoreactive bands were assigned based on their expected molecular weights and the predicted sizes of reporter-derived proteins. The full-length protein was defined as the band corresponding to the expected size of the GFP–TPI or TPI–GFP fusion protein, depending on the construct design. Truncated proteins were identified as lower molecular weight species consistent with premature translation termination at the introduced PTC. In PTC-containing constructs, readthrough-derived full-length products were detected at the position corresponding to the intact fusion protein, indicating translational readthrough of the PTC. Reinitiation-derived proteins were identified as distinct GFP-sized bands (∼26 kDa) or as bands corresponding to products initiated at downstream AUG codons (positions 14, 109, or 217) in engineered constructs.

Band intensities were quantified by densitometric analysis using Fiji (ImageJ, version 1.2.3). Integrated intensities were measured after local background subtraction and normalized to the corresponding loading control within the same lane. Quantification was performed separately for each protein species. All Western blot experiments were conducted with at least three independent biological replicates to ensure reproducibility of protein isoform detection and relative abundance measurements.

#### RNA extraction and RT-qPCR

Cells were transfected and harvested at the indicated time points, washed twice with ice-cold PBS, and lysed directly in culture dishes using the SuperPrep™ II Cell Lysis & RT Kit for qPCR (Toyobo, Osaka, Japan) according to the manufacturer’s instructions. Total RNA was treated with DNase I to remove genomic DNA contamination and quantified using a NanoDrop One spectrophotometer (Thermo Fisher Scientific). RNA quality was assessed prior to downstream analysis, and samples were stored at −80°C until use.

First-strand cDNA was synthesized from 0.5-1 μg of total RNA using SuperScript IV Reverse Transcriptase (Invitrogen, Thermo Fisher Scientific) according to the manufacturer’s instructions. Resulting cDNA samples were diluted with nuclease-free water prior to quantitative PCR. Quantitative PCR was performed using PowerTrack SYBR Green Master Mix (Applied Biosystems, Thermo Fisher Scientific) on a real-time PCR system. Reactions were performed in technical duplicates for each biological replicate.

At the end of amplification, melt curve analysis was conducted to confirm amplification specificity and absence of primer–dimer formation. Reporter mRNA abundance was quantified using primers specific to the expressed reporter transcript. For the 5FP-TPI constructs, primers were designed to amplify the EGFP-TPI junction. The mCherry-TPI control transcript was quantified separately using primer targeting mCherry-TPI junction. Primer pairs were designed to span exon-exon junctions. This enabled amplification of spliced reporter-derived cDNA and reduced the possibility of amplifying endogenous TPI transcripts or residual plasmid DNA.

Relative transcript levels were calculated using the 2^−ΔΔCt method and normalized to the corresponding wild-type (WW) control. All experiments were performed with at least three independent biological replicates. Data are presented as mean ± SD unless otherwise indicated. Primer sequences are listed in Supplementary Table 2.

## Supporting information

Supplementary Materials (DOCX)

## Acknowledgments

We thank members of the Singer laboratories for discussions. This work was supported by the World Premier International Research Center Initiative (WPI), MEXT, Japan; the Naito Grant for Female Scientists; the Mitani Foundation for Research Grant to HS.

## Author contributions

Conceptualization: HS

Methodology: HS

Investigation: NJP

Formal analysis: NJP, HS

Visualization: NJP, HS

Funding acquisition: HS

Project administration: HS

Resources: HS

Supervision: HS

Writing – original draft: NJP

Writing – review & editing: HS

## Data and materials availability

All data and code necessary to evaluate and reproduce the results reported in this paper are available in the supplementary materials. Plasmids, engineered cell lines, and other materials generated during this study are available from the corresponding author upon reasonable request.

