## Supplementary Materials (DOCX) for "Position-Dependent NMD Generates Diverse Protein Outcomes"

**This PDF file includes:**

Supplementary Text

Figs. S1 to S4

Supplementary Table 1 to 3

**Other Supplementary Materials for this manuscript include the following:**

R script code

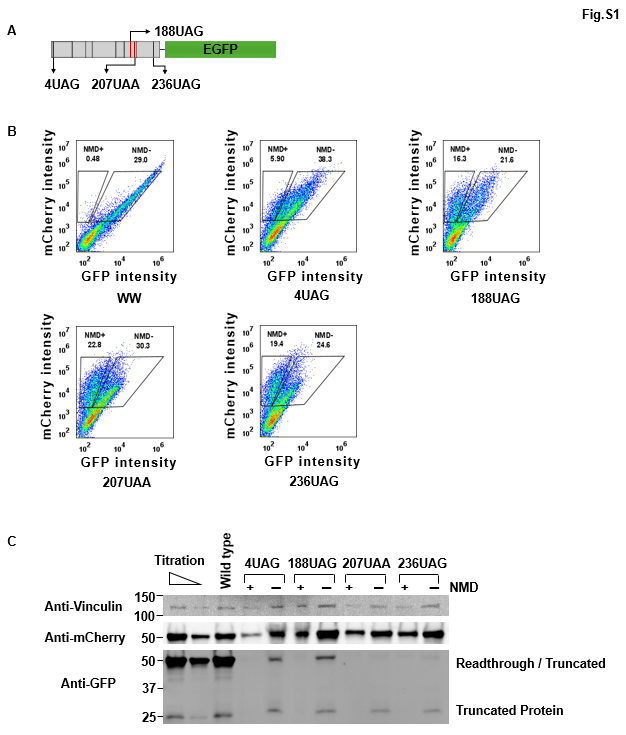
**Supplement. 1| 3’GFP tagging constructs not only detect Readthrough but also reinitiation.**

(A) A schematic diagram of TPI-GFP (3FP) constructs, where GFP inserted after TPI gene.
(B) Flow cytometry of TPI-GFP (3FP) constructs shows GFP fluorescence intensities in the PTC mutant.
(C) Western blot detects truncated TPI fusion protein of the GFP sized and shows readthrough at 5UAG.

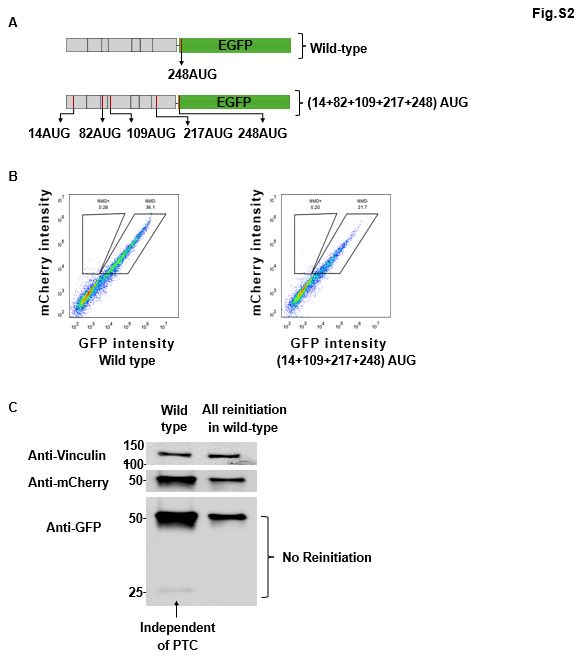

**Supplement. 2| Introducing multiple secondary start codon in wild-type for translation reinitiation**

(A) Schematic of TPI–GFP (3FP) constructs containing multiple AUG initiation codons in wild-type at different position.
(B) Flow cytometry analysis of TPI–GFP (3FP) constructs, showing EGFP and mCherry fluorescence distributions at the single-cell level.
(C) Western blot analysis of TPI–GFP (3FP) constructs showing no protein outputs generated from multiple AUG codons in wild-type constructs.

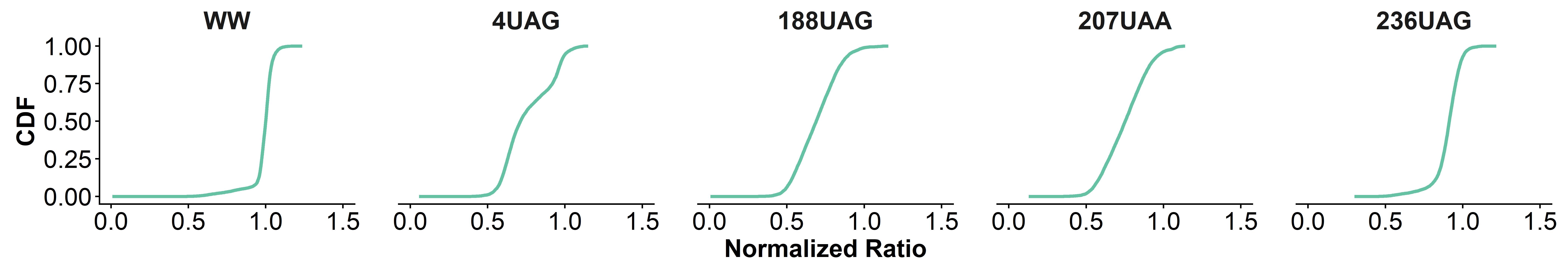

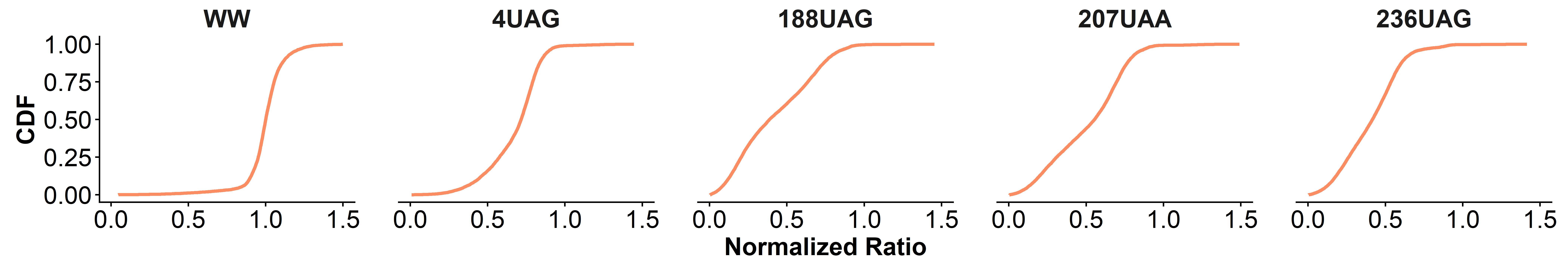

**Fig.S3**

**Supplement. 3| Cumulative distribution functions (CDFs) of normalized reporter expression.**
CDFs of normalized EGFP/mCherry ratios were generated for each nonsense mutation condition for 5FP (top) and 3FP (bottom) reporters. For each cell, EGFP intensity was normalized to mCherry and further normalized to the median EGFP/mCherry ratio of the wild-type (WW) control within the same experiment.

The CDFs represent the cumulative fraction of cells as a function of normalized expression, allowing comparison of distribution shape and tail behavior across conditions. These analyses reveal shifts in the proportion of cells with higher or lower NMD efficiency and highlight cell-to-cell heterogeneity that is not captured by median-based statistics.

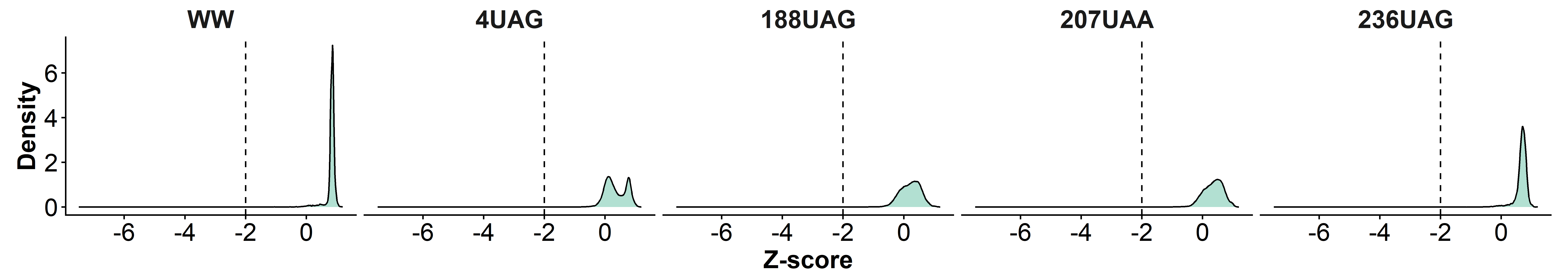

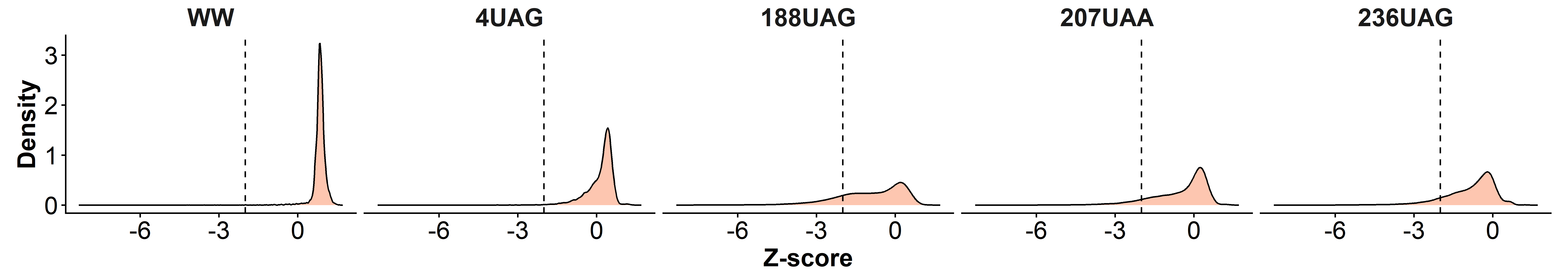

**Fig.S4**

**Supplement. 4| Density plots of Z-scores derived from log₁₀-transformed, normalized EGFP/mCherry ratios are shown for each condition.**

Z-scores were calculated across the dataset to represent the relative deviation of individual cells from the population mean.

Dashed lines indicate a Z-score threshold of −2, highlighting cells with markedly reduced reporter expression consistent with strong NMD activity. Separate panels show distributions for each condition for 5FP (top) and 3FP (bottom) reporters.

These distributions reveal shifts in population structure and the relative enrichment of cells with low expression, providing an additional measure of NMD-associated heterogeneity.

Supplementary Table 1. Pairwise statistical comparisons of reporter activity

| Type | group1 | group2 | n1 | n2 | statistic | p | p.adj | p.adj.signif |
| --- | --- | --- | --- | --- | --- | --- | --- | --- |
| 3FP | 188UAG | 207UAA | 13087 | 12277 | 63373294 | 3.12E-186 | 3E-186 | **** |
| 3FP | 188UAG | 236UAG | 13087 | 15278 | 1.01E+08 | 0.047 | 0.047 | * |
| 3FP | 188UAG | 4UAG | 13087 | 11570 | 31703142 | 0 | 0 | **** |
| 3FP | 188UAG | WW | 13087 | 6305 | 1518871 | 0 | 0 | **** |
| 3FP | 207UAA | 236UAG | 12277 | 15278 | 1.2E+08 | 0 | 0 | **** |
| 3FP | 207UAA | 4UAG | 12277 | 11570 | 40616281 | 0 | 0 | **** |
| 3FP | 207UAA | WW | 12277 | 6305 | 2000910 | 0 | 0 | **** |
| 3FP | 236UAG | 4UAG | 15278 | 11570 | 25145257 | 0 | 0 | **** |
| 3FP | 236UAG | WW | 15278 | 6305 | 1256590 | 0 | 0 | **** |
| 3FP | 4UAG | WW | 11570 | 6305 | 3049824 | 0 | 0 | **** |
| 5FP | 188UAG | 207UAA | 8845 | 9975 | 33342419 | 2.26E-184 | 2.8E-184 | **** |
| 5FP | 188UAG | 236UAG | 8845 | 10593 | 8662605 | 0 | 0 | **** |
| 5FP | 188UAG | 4UAG | 8845 | 8485 | 29399960 | 1.80E-134 | 2.0E-134 | **** |
| 5FP | 188UAG | WW | 8845 | 12149 | 3945655 | 0 | 0 | **** |
| 5FP | 207UAA | 236UAG | 9975 | 10593 | 18402202 | 0 | 0 | **** |
| 5FP | 207UAA | 4UAG | 9975 | 8485 | 42418803 | 0.782 | 0.782 | ns |
| 5FP | 207UAA | WW | 9975 | 12149 | 8684291 | 0 | 0 | **** |
| 5FP | 236UAG | 4UAG | 10593 | 8485 | 67222230 | 0 | 0 | **** |
| 5FP | 236UAG | WW | 10593 | 12149 | 18799656 | 0 | 0 | **** |
| 5FP | 4UAG | WW | 8485 | 12149 | 10105166 | 0 | 0 | **** |

Supplementary Table 2. Pairwise statistical comparisons of reporter activity of reinitiation

| group1 | group2 | n1 | n2 | p | p.adj | p.adj.signif | g1_Median_  Norm_Ratio | g1_SD_Log_Ratio | g2_Median_Norm_Ratio | g2_SD_Log_Ratio |
| --- | --- | --- | --- | --- | --- | --- | --- | --- | --- | --- |
| WW | (14+82)AUG | 3343 | 5876 | 0 | 0 | **** | 1 | 0.05578 | 0.753043 | 0.071048 |
| WW | (82+109)AUG | 3343 | 3777 | 0 | 0 | **** | 1 | 0.05578 | 0.71791 | 0.085401 |
| WW | (82+217)AUG | 3343 | 4803 | 0 | 0 | **** | 1 | 0.05578 | 0.718812 | 0.069699 |
| WW | (14+82+248)AUG | 3343 | 10125 | 0 | 0 | **** | 1 | 0.05578 | 0.758043 | 0.077064 |
| WW | (14+82+109+217+248)AUG | 3343 | 4932 | 0 | 0 | **** | 1 | 0.05578 | 0.763367 | 0.072485 |
| (14+82)AUG | (82+109)AUG | 5876 | 3777 | 4.03E-13 | 5.04E-13 | **** | 0.753043 | 0.071048 | 0.71791 | 0.085401 |
| (14+82)AUG | (82+217)AUG | 5876 | 4803 | 3.75E-15 | 5.62E-15 | **** | 0.753043 | 0.071048 | 0.718812 | 0.069699 |
| (14+82)AUG | (14+82+248)AUG | 5876 | 10125 | 0.95 | 0.95 | ns | 0.753043 | 0.071048 | 0.758043 | 0.077064 |
| (14+82)AUG | (14+82+109+217+248)AUG | 5876 | 4932 | 3.58E-16 | 5.97E-16 | **** | 0.753043 | 0.071048 | 0.763367 | 0.072485 |
| (82+109)AUG | (82+217)AUG | 3777 | 4803 | 0.196 | 0.21 | ns | 0.71791 | 0.085401 | 0.718812 | 0.069699 |
| (82+109)AUG | (14+82+248)AUG | 3777 | 10125 | 2.85E-12 | 3.29E-12 | **** | 0.71791 | 0.085401 | 0.758043 | 0.077064 |
| (82+109)AUG | (14+82+109+217+248)AUG | 3777 | 4932 | 4.53E-41 | 9.71E-41 | **** | 0.71791 | 0.085401 | 0.763367 | 0.072485 |
| (82+217)AUG | (14+82+248)AUG | 4803 | 10125 | 6.43E-14 | 8.77E-14 | **** | 0.718812 | 0.069699 | 0.758043 | 0.077064 |
| (82+217)AUG | (14+82+109+217+248)AUG | 4803 | 4932 | 2.84E-48 | 7.10E-48 | **** | 0.718812 | 0.069699 | 0.763367 | 0.072485 |
| (14+82+248)AUG | (14+82+109+217+248)AUG | 10125 | 4932 | 7.21E-21 | 1.35E-20 | **** | 0.758043 | 0.077064 | 0.763367 | 0.072485 |

Supplementary Table 3. List of Primers

| **Primer Name** | **Sequence (5' → 3')** |
| --- | --- |
| 3FP_PonA_NheI_TPI_F | GAACTCAGACACCATACTGCGGCTAGCATGGCGCCCTCCAGGAAGTTCTTCG |
| 3FP_TPI_KpnI_GFP_R | GCTCCTCGCCCTTGCTCACCATAATCGAGGTACCTTGTTTGGCATTGATGATGTCC |
| 3FP-MluI-mCherry TPI_F | GGGATCTGAGCACGCGAGCTACGCGTTCATTGTTTGGCATTGATGATG |
| 3FP-XhoI-TPI_R | GAACTCAGACACGAGCTCCTCGAGGCGATGGCGCCCTCCAGGAAGTTCTTC |
| 5FP-GFP-NotI-TPI_F | GGACGAGCTGTACAAGTACCGATGCGGCCGCTCTAGCATGGCGCCCTCCAGGAAGTTC |
| 5FP- SpeI-TPI_R | GGTATGGCTGATTATGATCACTAGTCGTATGGTTTTCGGATCGACGAAAGTCATTGTTTGGCATTGATG |
| 5FP-mCherry-XhoI-TPI_F | GACGAGCTGTACAAGCACGAGCTCCTCGAGGAATTCATGGCGCCCTCCAGGAAGTTCTTC |
| 5FP- ClaI-TPI_R | CCTCTGGAGATATCGTCGACAAGCTTATCGATTCATTGTTTGGCATTGATG |
| 3FP-GFP-F | ATGGTGAGCAAGGGCGAGGAGCTGTTCACCGGGG |
| 3FP_TPI_BamHI_GFP_R | GCTCCTCGCCCTTGCTCACCATAATCGAGGATCCTTGTTTGGCATTGATGATGTCC |
| 3FP-TPI-Ex6-189M-F | GGAAGTACACGAGAAGCTCTGAGGATGGCTGAAGTCC |
| 3FP-TPI-Ex6-189M-R | TCAGCCATCCTCAGAGCTTCTCGTGTACTTCC |
| 3FP-TPI-Ex6-208M-F | GCTCAGAGCACCCGTATCATTTAAGGAGGTGAGTGGCTTTGGTTCCCG |
| 3FP-TPI-Ex6-208M-R | GGAACCAAAGCCACTCACCTCCTTAAATGATACGGGTGCTCTGAGCCACC |
| TPI-Ex6-189(UAG)PTC-F | GTACACGAGAAGCTCTAGGGATGGCTGAAG |
| TPI-Ex6-189(UAG)PTC-R | CTTCAGCCATCCCTAGAGCTTCTCGTGTAC |
| TPI-Ex6-189(UAA)PTC-R | CTTCAGCCATCCTTAGAGCTTCTCGTGTAC |
| TPI-Ex6-189(UAA)PTC-F | GTACACGAGAAGCTCTAAGGATGGCTGAAG |
| TPI-Ex7-237(PTC)-F | GCTTCCCTCTAGCCCGAATTC |
| TPI-Ex7-237(PTC)-R | GAATTCGGGCTAGAGGGAAGC |
| 3FP-TPI-Ex7-237(PTC)-BamHI-R | GCTCACCATAATCGAGGATCCTTGTTTGGCATTGATGATGTCCACGAATTCGGGCTAGAGGGAAG |
| 3FP-TPI-Ex1-5(PTC)-NheI-F | CAGACACCATACTGCGGCTAGCATGGCGCCCTCCAGGTAGTTCTTCGTTG |
| 5FP-TPI-Ex1-5(PTC)-NotI-F | GCTGTACAAGTACCGATGCGGCCGCTCTAGCATGGCGCCCTCCAGGTAGTTCTTCGTTG |
| TPI-Ex7-Linker(ttg)-GFP-F | CAATGCCAAACAAGGATCCTCGATTTTGGTGAGCAAGGGCGAGGAGCTG |
| TPI-Ex7-Linker(ttg)-GFP-R | GGTGAACAGCTCCTCGCCCTTGCTCACCAAAATCGAGGATCCTTGTTTG |
| 3FP-NheI-Flag_tag-TPI-F | CTCAGACACCATACTGCGGCTAGCATGGACTACAAGGACGACGACGACAAGGCGCCCTCCAGGAAGTTC |
| 3FP-NheI-Flag_tag-TPI(5UAG)-F | CTCAGACACCATACTGCGGCTAGCATGGACTACAAGGACGACGACGACAAGGCGCCCTCCAGGTAGTTC |
| 5FP-TPI-Ex7-Flag-SpeI-R | GGTATGGCTGATTATGATCACTAGTCGTATGGTTTTCGGATCGACGAAAGTCACTTGTCGTCGTCGTCCTTGTAGTCTTGTTTGGCATTGATGATG |
| 3FP-TPI-Ex1-4PTC-NheI-14TTG-F | CAGACACCATACTGCGGCTAGCATGGCGCCCTCCAGGTAGTTCTTCGTTGGGGGAAACTGGAAGTTGAACGGGCGGAAGCAGAGT |
| 3FP-TPI-Ex4-109ATG-F | CAAGTCTGTTTCTCAACAGCTGATGGGGCAGAAAGTGGCCCATG |
| 3FP-TPI-Ex4-109ATG-R | CATGGGCCACTTTCTGCCCCATCAGCTGTTGAGAAACAGACTTG |
| 3FP-TPI-Ex7-217ATG-F | CTCTGTGACTGGGGCAACCTGCATGGAGCTGGCCAGCCAGCCTGATGTG |
| 3FP-TPI-Ex7-217ATG-R | CACATCAGGCTGGCTGGCCAGCTCCATGCAGGTTGCCCCAGTCACAGAG |
| GFP-3'-F | AGCGCGATCACATGGTCCTGCT |
| qPCR-GFP-110-F | ATGCCACCTACGGCAAGC |
| qPCR-GFP-215-R | AAGCACTGCACGCCGTAG |
| qPCR-mCherry-F | TTGGTCACCTTCAGCTTG |
| qPCR-mCherry-R | GAGTTCATGCGCTTCAAG |
| mCherry-3end-qPCR-F | ATCGTGGAACAGTACGAA |
| TPI-Ex1-Ex2-qPCR-R | ACAAACCACCTCGGTGTC |
